## Supplemental Data for "Serine Phosphorylation Regulates the P-type Potassium pump KdpFABC"

### Supplemental Figure Legends

Suppl. Figure 1. Detection of Ser162 phosphorylation by mass spectrometry. (A) MS/MS spectrum of phosphopeptide from KdpFABC with m/z 705.3376 and sequence SAITGE$^{162}$pSAPVIRE. The observed fragment ions were used to confirm that Ser162 is the site of phosphorylation. B) Predicted MS/MS fragment ions for the phosphopeptide SAITGE$^{162}$pSAPVIRE. Fragment ions in red were detected in the spectrum in panel A. The data analysis was generated by the Mascot search engine which reported a 99.7% probability that the site of phosphorylation was on Ser162 and a combined probability of 0.3% for Ser156 and Thr159.

Suppl. Figure 2. Preparation of KdpFABC and analysis of serine phosphorylation using Phos-tag stain. (A) Coomassie stained SDS-PAGE of purified preparations used for EP measurements. Although KdpA, KdpB and KdpC are readily visible, KdpF has run off the bottom of the gel due to its small mass. (B) Elution profile from the size-exclusion column shows that the purified protein is monodisperse. (C) ATPase activities (mean and SEM) of preparations shown in panel A before and after LPP treatment to remove serine phosphorylation. Triplicate measurements were made from each preparation. (D) Phos-tag staining of various preparations of KdpFABC before and after $K^+$ shock for the indicated periods of time. KdpB has varying levels of Ser162 phosphorylation, which has been quantified and plotted in Fig. 4. Unexpectedly, we also observed a constant level of staining of KdpA, which could reflect a tightly bound lipid. This idea is supported by data in panel E, which shows that LPP treatment reduced the staining associated with KdpB, but did not affect the staining of KdpA.

Suppl. Figure S3. Steady-state levels of aspartyl phosphate (EP) are constant at 4°C. (A) Individual experiments comparing time-dependence of EP formation in the absence of $K^+$. (B) Data from the same constructs in the presence of 10 mM $K^+$. EP formation was initiated by addition of [$\gamma^{32-}$P]-ATP and aliquots were taken at various time intervals and quenched with TCA. Data was collected in triplicate and the specific constructs are indicated in the legend.

Suppl. Figure S4. Electrogenic transport of $K^+$ by KdpFABC. Transport was initiated by addition of ATP (black arrow), which generates a decrease in fluorescence as the membrane voltage builds up. (A, C, E) Raw data from DiSC3 fluorescence generated by reconstituted proteoliposomes. In panel A, the red arrow indicate an addition of valinomycin, which abolished the membrane voltage. (B, D, F) Analysis of these data to determine the initial rate, which involved normalizing the signal, determining the rate of quenching and fitting the curve with an exponential. The exponential fit is indicated as a red line, the quench rate as a blue line and the initial slope as a green line. These data come from WT protein subjected to $K^+$ shock before (A,B) and after (C,D) LPP treatment as well as from the D307A mutant (E,F), which serves as a negative control (indicated at the bottom of each plot). (G) Comparison of transport rates from five different preparations based on the initial slopes from three independent experiments. Inhibition of transport by $K^+$-shock and recovery with LPP treatment is consistent with results from ATPase activity measurements in Fig. 4, indicating that ATP hydrolysis and $K^+$ transport remain coupled in these preparations. Measurements were performed in triplicate and plotted as mean plus SEM.

Suppl. Figure S5. Interactions between the A-domain (yellow) and N-domain (red) mediated by serine phosphorylation. This interaction was seen in the X-ray structure, which was the starting point for MD simulations. Radial distribution functions and RMSD distributions indicate that this interaction was stable in the presence of serine phosphorylation over the course of the 200 ns simulations, but not in its absence. (A) Radial distribution functions for center of masses from side chains of Glu161 and Lys357. These data represent 2 $\mu$s of simulation trajectories (concatenation of ten individual 200 ns simulations). (B) Radial distribution functions for center of masses from side chains of Ser162 and Arg357. (C) Root mean square deviations (RMSD) for side chains of Glu161 and Lys357. (D) RMSD for side chains of Ser162 and Arg357. RMSD was calculated using all atoms from each residue from the last 100 ns of trajectories after correcting for bulk translational motion. (E) Representative snapshot from a PHOS simulation showing the proximity of Glu161 and phosphorylated Ser162 to Lys357 and Arg363. (F) Representative snapshot from a SER simulation showing that the interaction is disrupted in the absence of the phosphoserine. (G) Phos-tag staining of the K357A/R363A/Q116R mutant expressed in TK2498 shows robust serine phosphorylation of KdpB that is reduced by LPP treatment. (H) ATPase activities from the K357A/R363A/Q116R mutant show an ~4-fold stimulation resulting from LPP treatment, thus producing ATPase levels comparable to WT and indicating that the interaction between A- and N-domains is not essential for inhibition of KdpFABC.

Suppl. Figure S6. Distribution of eigenvalues from the covariance matrix of atomic fluctuations from PHOS (blue) and SER (orange) simulations. The first two eigenvectors represent the largest deviations in the dataset and have been used in Fig. 8 to analyze the global collective motions.

Suppl. Figure S7. Water accessibility of the canonical substrate binding site in KdpB during the simulations. (A) Radial distribution functions calculated for water molecules relative to the center of mass for Pro264, Thr265, Asp583 and Lys586 from KdpB, which surround the canonical substrate binding site. These traces represent the aggregate of all ten simulations for phosphorylated and unphosphorylated KdpFABC. The higher peak for PHOS simulations indicate a much greater penetration by water. (B) Initial configuration of the canonical site for both the SER and PHOS simulations with a water molecule (blue) bound at the canonical ion binding site as seen in the X-ray structure. (C) and (D) Final frames from PHOS and SER simulations respectively. The canonical substrate binding site is significantly more hydrated in the PHOS simulations with water penetrating the site from the cytoplasmic side of the membrane (bottom of the panel).

### Suppl. Table 1. KdpFABC mutants

| annotation | mutations | strain | promoter | condition | Figures |
|---|---|---|---|---|---|
| WT | - | TK2499 | KdpE | μM $K^+$ | 1,2,3 |
| KdpA-E370A | KdpA-E370A | TK2498 | KdpE | 0.2 mM $K^+$ | 1 |
| KdpA-R400A/T401A | KdpA-R400A KdpAT401A | TK2498 | KdpE | 0.2 mM $K^+$ | 1 |
| KdpA-S517A | KdpA-S517A | TK2498 | KdpE | 0.2 mM $K^+$ | 1 |
| KdpA-R400A | KdpA-R400A | TK2498 | KdpE | 0.2 mM $K^+$ | 1 |
| KdpA-R493A | KdpA-R493A | TK2498 | KdpE | 0.2 mM $K^+$ | 1 |
| KdpB-D300A | KdpB-D300A | TK2498 | KdpE | 0.2 mM $K^+$ | 1 |
| KdpB-D302 | KdpB-D302 | TK2498 | KdpE | 0.2 mM $K^+$ | 1 |
| KdpB-D300A/D302A | KdpB-D300A/ KdpB-D302A | TK2498 | KdpE | 0.2 mM $K^+$ | 1 |
| KdpB-D307A | KdpB-D307A | TK2498 | KdpE | 0.2 mM $K^+$ | 1 |
| KdpB-D583A | KdpB-D583A | TK2498 | KdpE | 0.2 mM $K^+$ | 1 |
| KdpB-K586A | KdpB-K586A | TK2498 | KdpE | 0.2 mM $K^+$ | 1 |
| K357/R363/ Q116R | KdpA-Q116R/ KdpB-K357/ KdpB-R363 | TK2498 | KdpE | 0.2 mM $K^+$ | S5 |
| Q116R | KdpA-Q116R | Top10 | pBAD | LB media* | 3,4 |
| S162A/Q116R | KdpA-Q116R/ KdpB-S162A | Top10 | pBAD | LB media* | 4 |
| S162D/Q116R | KdpA-Q116R/ KdpB-S162D | Top10 | pBAD | LB media* | 4 |
| D307A/Q116R | KdpA-Q116R/ KdpB-D307A | Top10 | pBAD | LB media* | 4 |
| S162A/E161Q/ Q116R | KdpA-Q116R/ KdpB-S162A/ KdpB-E161Q | Top10 | pBAD | LB media* | 4 |

* LB media has been reported to contain 8 mM $K^+$ (Su et al., 2009)

A

E.SAITGE162pSAPVIRE.S.
Observed m/z: 705.3376, Experimental mass: 1408.6606, Calculated mass: 1408.6599

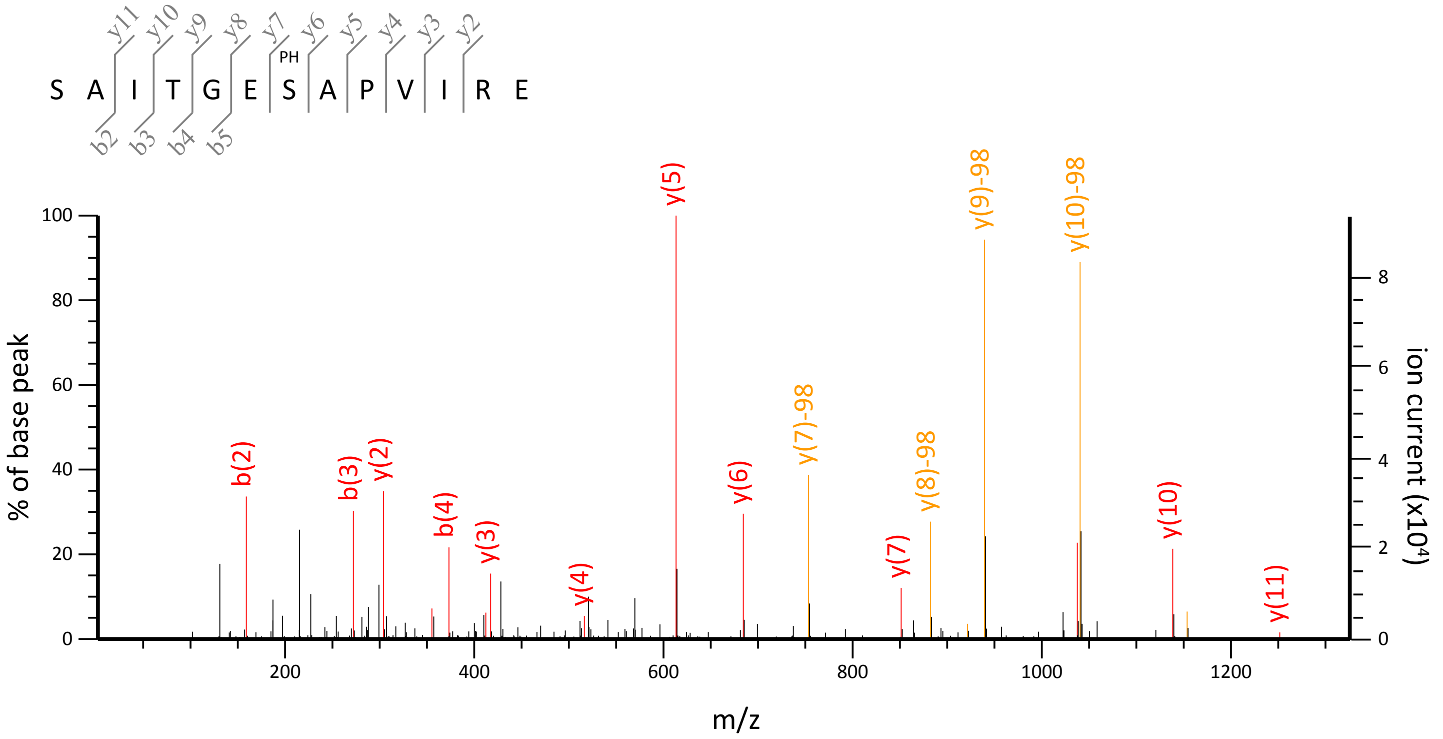


B

| # | b | b++ | b* | b*++ | b0 | b0++ | Seq | y | y++ | y* | y*++ | y0 | y0++ | # |
|---|---|---|---|---|---|---|---|---|---|---|---|---|---|---|
| 1 | 88.0393 | 44.5233 | | | 70.0287 | 35.5180 | **S** | | | | | | | 13 |
| 2 | **159.0764** | 80.0418 | | | 141.0659 | 71.0366 | **A** | 1224.6583 | 612.8328 | 1207.6317 | 604.3195 | 1206.6477 | 603.8275 | 12 |
| 3 | **272.1605** | 136.5839 | | | 254.1499 | 127.5786 | **I** | **1153.6212** | 577.3142 | 1136.5946 | 568.8009 | 1135.6106 | 568.3089 | 11 |
| 4 | **373.2082** | **187.1077** | | | **355.1976** | 178.1024 | **T** | **1040.5371** | **520.7722** | 1023.5106 | 512.2589 | 1022.5265 | 511.7669 | 10 |
| 5 | 430.2296 | 215.6185 | | | **412.2191** | 206.6132 | **G** | **939.4894** | 470.2483 | 922.4629 | 461.7351 | 921.4789 | 461.2431 | 9 |
| 6 | 559.2722 | 280.1397 | | | 541.2617 | 271.1345 | **E** | **882.4680** | 441.7376 | 865.4414 | 433.2243 | 864.4574 | 432.7323 | 8 |
| 7 | 628.2937 | 314.6505 | | | 610.2831 | 305.6452 | **S** | **753.4254** | 377.2163 | 736.3988 | 368.7030 | 735.4148 | 368.2110 | 7 |
| 8 | 699.3308 | 350.1690 | | | 681.3202 | 341.1638 | **A** | **684.4039** | 342.7056 | 667.3774 | 334.1923 | 666.3933 | 333.7003 | 6 |
| 9 | 796.3836 | 398.6954 | | | 778.3730 | 389.6901 | **P** | **613.3668** | 307.1870 | 596.3402 | 298.6738 | 595.3562 | 298.1817 | 5 |
| 10 | 895.4520 | 448.2296 | | | 877.4414 | 439.2243 | **V** | **516.3140** | 258.6606 | 499.2875 | 250.1474 | 498.3035 | 249.6554 | 4 |
| 11 | 1008.5360 | 504.7717 | | | 990.5255 | 495.7664 | **I** | **417.2456** | 209.1264 | 400.2191 | 200.6132 | 399.2350 | 200.1212 | 3 |
| 12 | 1164.6371 | 582.8222 | 1147.6106 | 574.3089 | 1146.6266 | 573.8169 | **R** | **304.1615** | 152.5844 | 287.1350 | 144.0711 | 286.1510 | 143.5791 | 2 |
| 13 | | | | | | | **E** | 148.0604 | 74.5339 | | | 130.0499 | 65.5286 | 1 |

Suppl. Figure 1

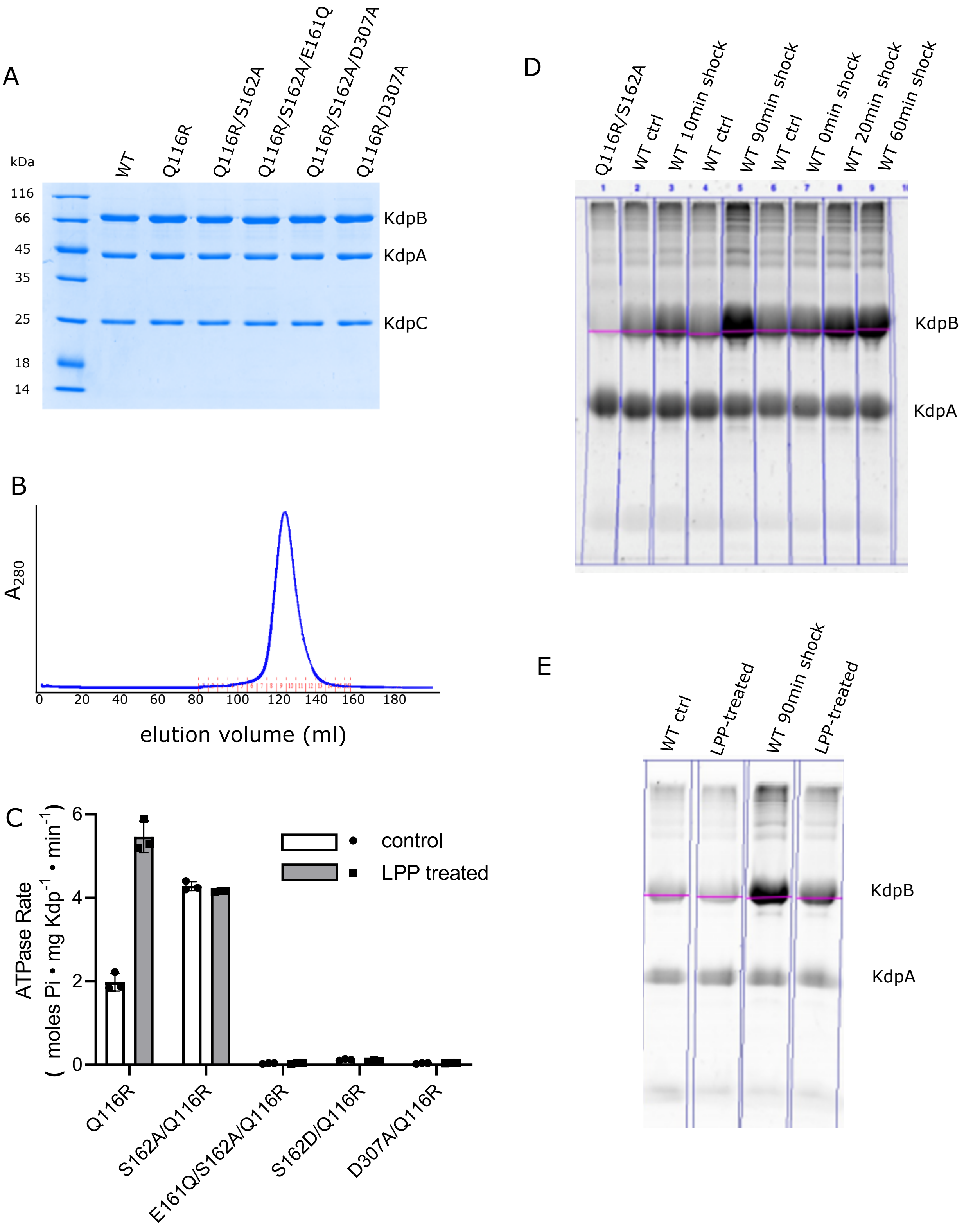


Suppl. Figure 2

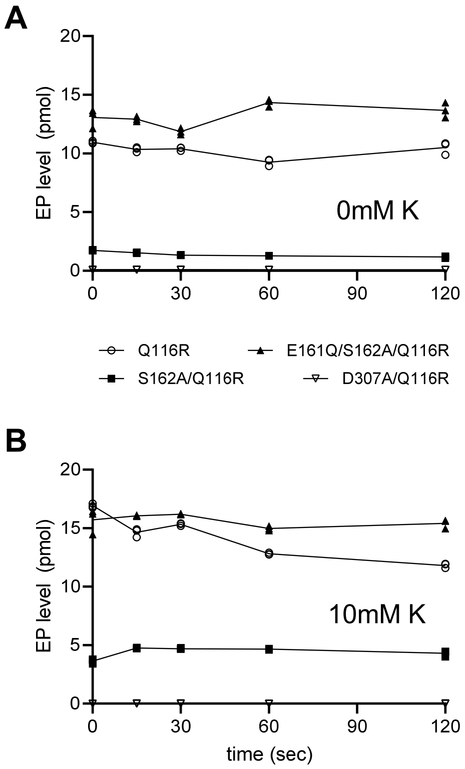


Suppl. Figure 3

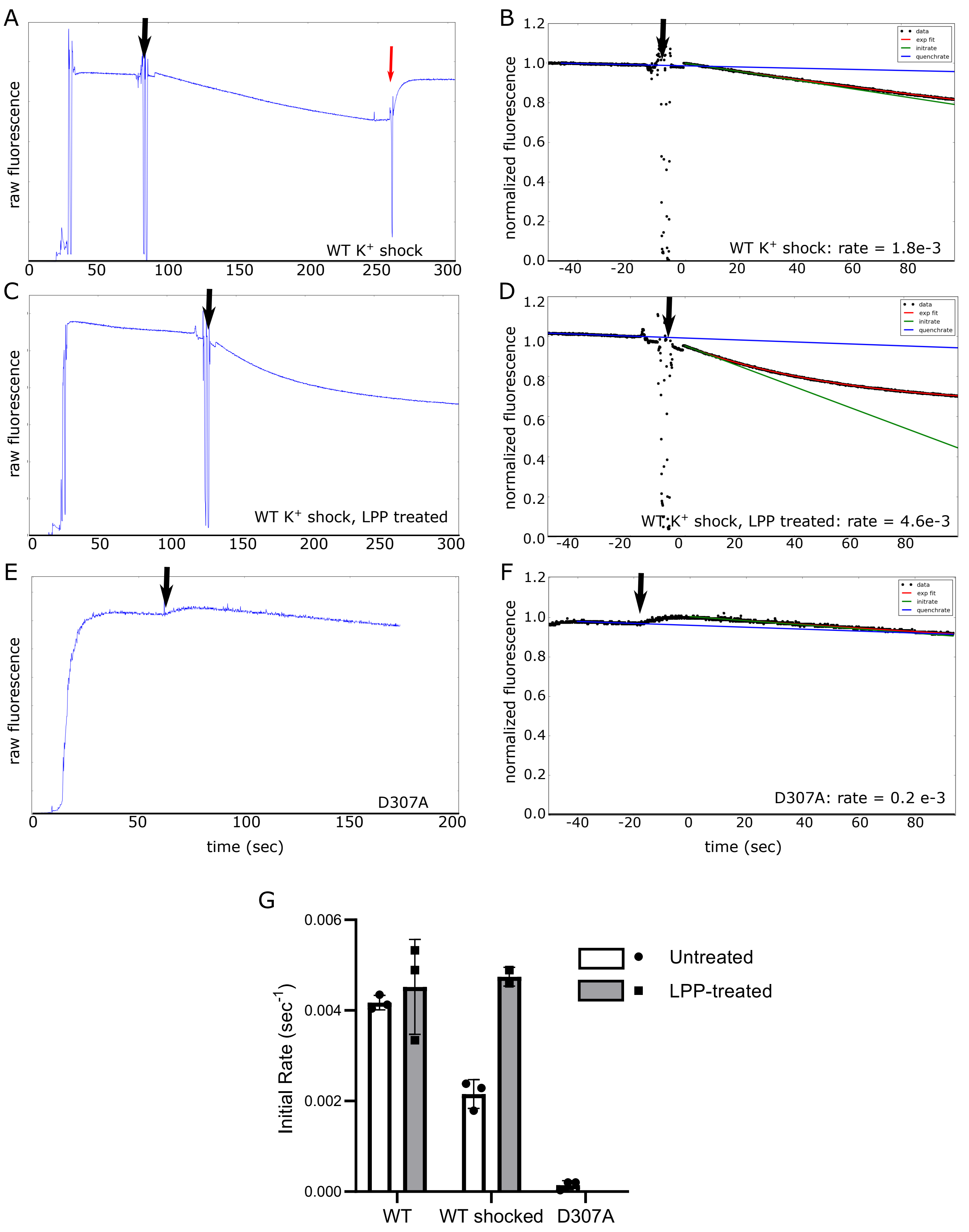


Suppl Figure 4

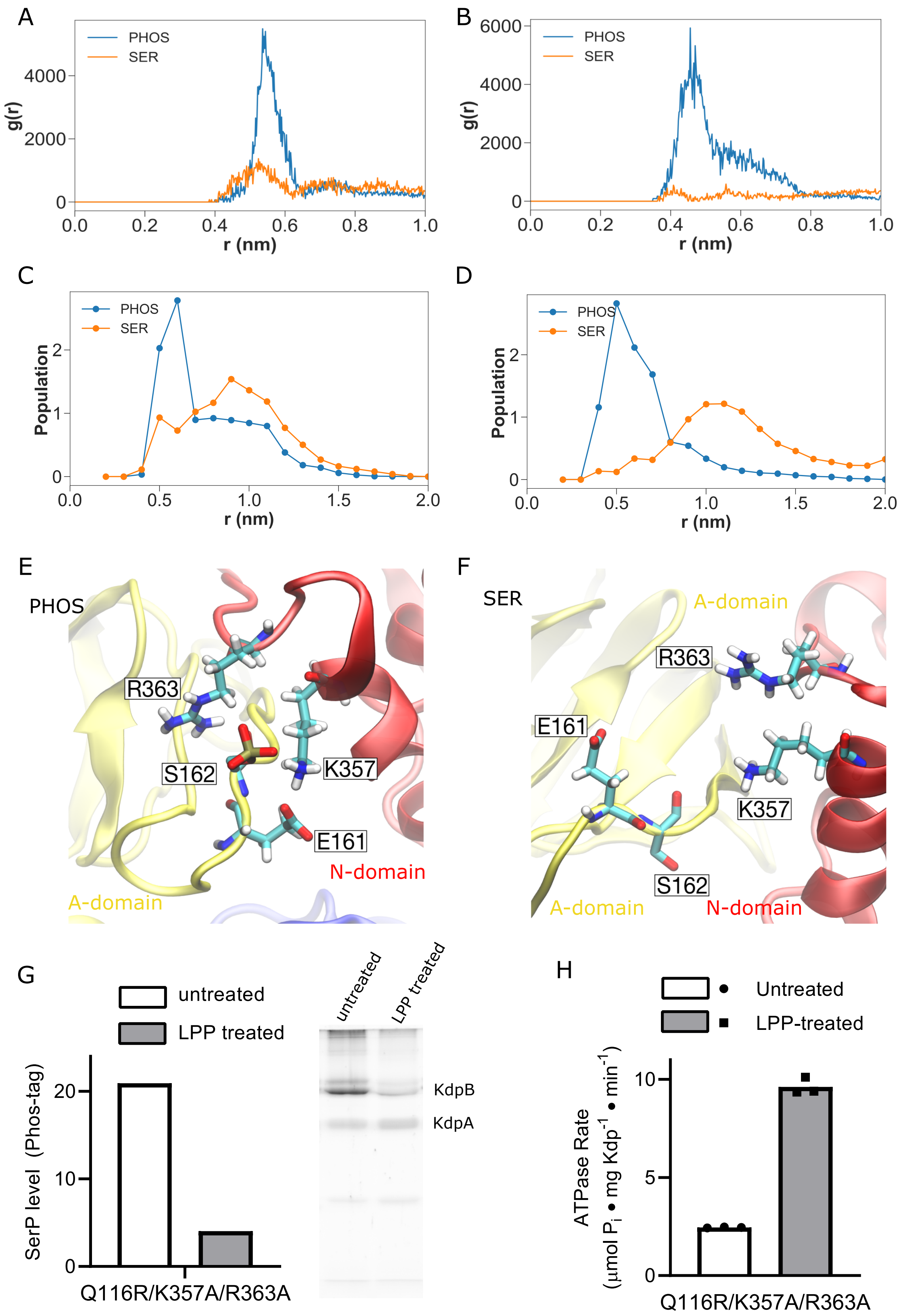


Suppl Figure 5

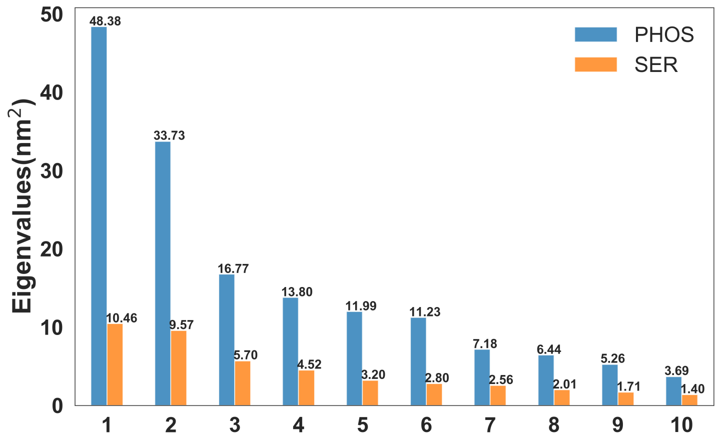


Suppl. Figure 6

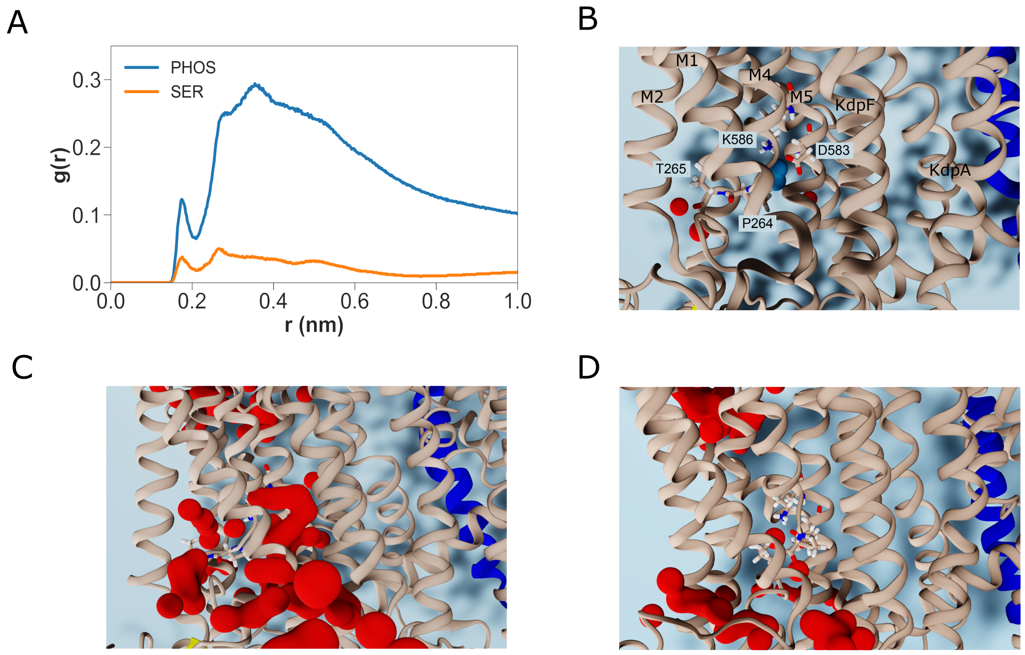


Suppl. Figure 7
